# Genome-wide characterization of the common bean kinome: catalog and insights into expression patterns and genetic organization

**DOI:** 10.1101/2022.08.05.503001

**Authors:** Alexandre Hild Aono, Ricardo José Gonzaga Pimenta, Caroline Marcela da Silva Dambroz, Francisco Cleilson Lopes Costa, Reginaldo Massanobu Kuroshu, Anete Pereira de Souza, Welison Andrade Pereira

**Affiliations:** Molecular Biology and Genetic Engineering Center (CBMEG), University of Campinas (UNICAMP), Campinas, Brazil; Department of Biology, Federal University of Lavras (UFLA), Lavras, Brazil; Instituto de Ciência e Tecnologia, Universidade Federal de São Paulo (UNIFESP), São José dos Campos, Brazil; Department of Plant Biology, Biology Institute, University of Campinas (UNICAMP), Campinas, Brazil

**Author notes:** Email addresses:* (Alexandre Hild Aono), (Ricardo José Gonzaga Pimenta), (Caroline Marcela da Silva Dambroz), (Francisco Cleilson Lopes Costa), (Reginaldo Massanobu Kuroshu), (Anete Pereira de Souza), (Welison Andrade Pereira).

**Keywords:** coexpression networks, duplication events, gene expression, kinase gene family, *Phaseolus vulgaris*, phylogenetic analyses

## Abstract

The protein kinase (PK) superfamily is one of the largest superfamilies in plants and is the core regulator of cellular signaling. Even considering this substantial importance, the kinome of common bean (*Phaseolus vulgaris*) has not been profiled yet. Here, we identified and characterised the complete set of kinases of common bean, performing an in-depth investigation with phylogenetic analyses and measurements of gene distribution, structural organization, protein properties, and expression patterns over a large set of RNA-Sequencing data. Being composed of 1,203 PKs distributed across all *P. vulgaris* chromosomes, this set represents 3.25% of all predicted proteins for the species. These PKs could be classified into 20 groups and 119 subfamilies, with a more pronounced abundance of subfamilies belonging to the receptor-like kinase (RLK)-Pelle group. In addition to provide a vast and rich reservoir of data, our study supplied insights into the compositional similarities between PK subfamilies, their evolutionary divergences, highly variable functional profile, structural diversity, and expression patterns, modeled with coexpression networks for investigating putative interactions associated with stress response.

## 1. Introduction

A kinome can be defined as an organism complete set of proteins that contain a kinase domain, which are denominated protein kinases (PKs). The kinase domain is characterized by a catalytic core consisting of 250 to 300 conserved amino acids with substrate specificity (Lehti-Shiu & Shiu, 2012; Wei et al., 2014). PKs have the ability to phosphorylate protein substrates, transferring a *γ*-phosphate residue from an ATP molecule to the hydroxyl group of a serine, threonine or tyrosine in the target protein (Hanks & Hunter, 1995; Liu et al., 2020). Through this process, PKs regulate the activity of their targets and, as a consequence, mediate diversified processes of an organism’s life, from its early development to its responses to biotic or abiotic stresses (Liu et al., 2015).

PKs are part of the largest and most conserved gene superfamily in plants (Liu et al., 2015), piquing the interest of researchers seeking to elucidate crucial mechanisms of plant vegetative and reproductive development, in addition to their responses to the environment. According to Lehti-Shiu & Shiu (2012), PKs from various plant species can be identified and classified into 115 families organized into several groups, including receptor-like kinase (RLK)-Pelle; cGMP-dependent protein kinase, and lipid signaling kinase families (AGC); calcium- and calmodulin-regulated kinase (CAMK); casein kinase 1 (CK1); cyclin-dependent kinase, mitogen-activated protein kinase, glycogen synthase kinase, and cyclin-dependent kinase-like kinase (CMGC); cyclic AMP-dependent protein kinase (cAPK); serine/threonine kinase (STE); and tyrosine kinase-like kinase (TKL). In fact, the functional classification of PKs based on the conservation and phylogeny of their catalytic domains enabled the first studies on these proteins (Liu et al., 2015). The first plant PK was isolated from pea (*Pisum sativum*) in 1973 (Keates, 1973), and the first plant PK DNA sequences were identified in common bean (*Phaseolus vulgaris*) and rice (*Oryza sativa*) using degenerate primers in 1989 (Lawton et al., 1989). Since then, the study of kinomes of different plant species at a genome-wide scale has been possible with the advancement of high-throughput sequencing technologies Liu et al. (2015).

Several studies have shown that plant kinomes are much larger than those of other eukaryotes (Liu et al., 2015). For instance, the human genome has 518 predicted kinases (Manning et al., 2002), while plant species have between 600 to 2,500 members (Lehti-Shiu & Shiu, 2012). This expansion in the repertoire of plant PKs can be attributed to frequent recent whole genome duplication (WGD) events associated with high rates of gene retention (Lehti-Shiu & Shiu, 2012). The increase in the complexity of a species can be associated with the expansion in the complexity of its proteins – which helps to explain the large variation in the number and diversity of PKs in different species. The study of kinomes demonstrates, based on PK representativeness, the relevance of this protein superfamily for the physiology of a species. As an illustration, 954 PKs were found in *Fragaria vesca* (Liu et al., 2020), 1,168 PKs in *Vitis vinifera* (Zhu et al., 2018b), 758 PKs in *Ananas comosus* (Zhu et al., 2018a), 2,168 PKs in *Glycine max* (Liu et al., 2015), 1,436 PKs in *Solanum lycopersicum* (Singh et al., 2014), 1,241 PKs in *Zea mays* (Wei et al., 2014), and 942 PKs in *Arabidopsis thaliana* (Zulawski et al., 2014).

Among the species of major interest in agriculture, the common bean (*P. vulgaris* L.) stands out with its fundamental importance for human consumption. Its grains are a source of protein, lysine, fiber (Messina, 2014), folate, and mineral salts, such as iron, zinc, magnesium and potassium (Mitchell et al., 2009). Additionally, they present resistant starch (Hutchins et al., 2012) and phenolic compounds with antioxidant potential (Marathe et al., 2011). Among many health benefits, beans act positively on cholesterol levels (Gunness & Gidley, 2010), blood glucose, inflammatory processes, metabolic syndrome and cardiovascular diseases (Messina, 2014). Given these attributes, bean production arises as a strategic action for agriculture, which justifies the maintenance of the large area destined for sowing this crop in the 2022 harvest (Conab, 2022).

Nevertheless, bean production is affected by several types of stresses, including fungal, viral and bacterial diseases (Basavaraja et al., 2020), insect and nematode pests (Singh & Schwartz, 2010), drought, and aluminium toxicity (Beebe et al., 2009). PKs have a well-established role in the response to both biotic and abiotic stresses, being part of highly complex signaling cascades (Ben Rejeb et al., 2014; Ye et al., 2017). Importantly, PK genes have been identified as the source of resistance to anthracnose, a devastating disease that can lead to yield losses of up to 100% in *P. vulgaris* (Pvu) (Melotto & Kelly, 2001; Oblessuc et al., 2015; Richard et al., 2021). Additional evidence of associations of these proteins with resistance to other diseases in common bean (Cooper et al., 2020; Vasconcellos et al., 2017) further highlight the importance of their characterization in this crop.

Given the potential of PKs for the development of plants and their interaction with the environment, it is essential that this superfamily is thoroughly described and analysed to enable biochemical and molecular inferences about various aspects of plant-environment interactions. Despite the availability of the common bean genome since 2014 (Schmutz et al., 2014), the kinome of Pvu has not yet been investigated. The aim of this study, therefore, was to identify, classify and catalogue the complete set of PKs of this species. Furthermore, phylogenetic analyses and predictions of chromosomal location and structural organization of genes encoding PKs were performed. Lastly, PK subfamilies had their gene expression estimated with a large set of RNA-Seq data, and genic interactions were modeled using coexpression networks.

## 2. Material and methods

### 2.1. Genome-wide kinase identification and phylogenetic analyses

Pvu gene, coding DNA, and protein-coding gene sequences were retrieved from the Pvu genome (v2.1) in Phytozome v.13 (Goodstein et al., 2012). For protein kinase (PK) identification, we selected hidden Markov models (HMMs) of typical kinase families from the Pfam database (El-Gebali et al., 2019): Pkinase (PH00069) and Pkinase Tyr (PF07714). All Pvu protein sequences were aligned against kinase HMMs using HMMER v.3.3 (Finn et al., 2011) with an E-value cut-off of 0.1 and a minimum domain coverage of 50% (Lehti-Shiu & Shiu, 2012). For genes with isoforms, only the longest variant was retained for further analyses.

Putative Pvu PKs were classified into subfamilies based on HMMs calculated with sequences from other 25 plant species (Lehti-Shiu & Shiu, 2012): *Aquilegia coerulea* (Aco), *Arabidopsis lyrata* (Aly), *Arabidopsis thaliana* (Ath), *Brachypodium distachyon* (Bdi), *Carica papaya* (Cpa), *Citrus clementina* (Ccl), *Citrus sinensis* (Csi), *Chlamydomonas reinhardtii* (Cre), *Cucumis sativus* (Csa), *Eucalyptus grandis* (Egr), *Glycine max* (Gma), *Manihot esculenta* (Mes), *Medicago truncatula* (Mtr), *Mimulus guttatus* (Mgu), *O. sativa* (Osa), *Populus trichocarpa* (Ptr), *Prunus persica* (Ppe), *Physcomitrella patens* (Ppa), *Ricinus communis* (Rco), *Selaginella moellendorffii* (Smo), *Setaria italica* (Sit), *Sorghum bicolor* (Sbi), *Vitis vinifera* (Vvi), *Volvox carteri* (Vca) and *Zea mays* (Zma).

To confirm the PKs’ subfamily classification, kinase domains from the set of identified PKs were aligned with Muscle v.3.8.31 (Edgar, 2004) and used for constructing a phylogenetic tree (1,000 bootstraps) with FastTreeMP v2.1.10 (Price et al., 2010) via CIPRES gateway (Miller et al., 2011). The generated tree visualization was assessed with the ggtree (Yu et al., 2017) and ggplot2 (Villanueva & Chen, 2019) packages on R statistical software (R Core Team, 2013).

### 2.2. Kinase characterization

The chromosomal location of Pvu PK genes was determined using the GFF file obtained from Phytozome, and visualized with MapChart v2.2 software (Voorrips, 2002). With this same file, we also estimated the gene organization of PK subfamilies through intron numbers. Several protein properties were evaluated for the Pvu kinome. The domain composition of PKs was characterized using the Pfam database and the HMMER web server (Finn et al., 2011); their subcellular localizations were predicted with the programs WoLF PSORT (Horton et al., 2007), CELLO v.2.5 (Yu et al., 2006) and LOCALIZER v.1.0.4 (Sperschneider et al., 2017). Transmembrane domains and N-terminal signal peptides were recognized with TMHMM v.2.0 Server (Krogh et al., 2001) and SignalP v.4.1 Server (Armenteros et al., 2019) respectively; and theoretical isoelectric points (pIs) and molecular weights predicted with the ExPASy server (Artimo et al., 2012). These properties were summarized with descriptive statistics and different plots constructed with the R statistical software (R Core Team, 2013). The functional annotation of the Pvu kinome was performed with the Blast2GO tool (Conesa & Götz, 2008) together with SWISS-PROT (Bairoch & Apweiler, 2000) and Uniprot (Consortium, 2019) databases. From the PK annotations, Gene Ontology (GO) terms (Ashburner et al., 2000) were retrieved and analysed via treemaps constructed using the REViGO tool (Supek et al., 2011).

### 2.3. Duplication events

The Multiple Collinearity Scan (MCScanX) toolkit (Wang et al., 2012) was used for identifying putative homologous PKs along the Pvu genome and categorizing duplication events, which were separated into tandem and segmental duplications and visualized using MapChart v2.2 (Voorrips, 2002) and Circos (Krzywinski et al., 2009) softwares, respectively. We also used MCScanX for calculating synonymous (Ks) and non-synonymous substitution (Ka) rates, and with Ks estimations, we calculated the date of duplication events using the formula *T* = *K*_*s*_*/*2*λ*, with *λ* representing the mean value of clock-like Ks rates (6.5 × 10^*−*9^) (Gaut et al., 1996).

### 2.4. RNA-Seq experiments and co-expression network modelling

PKs’ expression quantifications were assessed using RNA-Seq experiments detailed in Supplementary Table S1 and obtained from NCBI’s Sequence Read Archive (SRA) (Leinonen et al., 2010). We selected datasets containing samples from different bean genotypes (Negro Jamapa, SA118, SA36, Black Turtle Soup, G19833, Ispir, DOR364 and IAC-Imperador) and analysed in different tissues (leaves, stems, shoots, flowers, pods, seeds and nodules) (Hiz et al., 2014; Kamfwa et al., 2017; Khankhum et al., 2016; Lu et al., 2019; O’Rourke et al., 2014; Silva et al., 2019). RNA-Seq reads were downloaded and their quality assessed with FastQC software (Andrews, 2010). Using Trimmomatic v.0.39 (Bolger et al., 2014), we only retained reads with a minimum Phred score of 20 and larger than 30 bp, which were used for PK quantification using bean transcript sequences downloaded from Phytozome and the Salmon v.1.1.0 software (Patro et al., 2017) (k-mer of 17). All PK expression counts were normalized using transcripts per million (TPM) values. PK subfamilies’ expression was evaluated using a heatmap representation with the pheatmap R package (Kolde & Kolde, 2015), considering averaged TPM values and a complete-linkage hierarchical method with euclidean distances. Pairwise correlations between kinase subfamilies were calculated with Pearson correlation coefficients and used for modeling co-expression networks via igraph R package (Csardi et al., 2006). Each node in such a structure represents a kinase subfamily, and an edge a minimum correlation coefficient of 0.6. We created two different networks, separating RNA samples according to control and adverse experimental conditions. These networks were evaluated and compared considering: (i) their community structures assessed with a propagating label algorithm (Raghavan et al., 2007); (ii) hub scores calculated with Kleinberg’s hub centrality (Kleinberg, 1999); and (iii) edge betweenness measured with the number of geodesics passing through an edge (Brandes, 2001).

## 3. Results

### 3.1. Genome-wide identification and classification of common bean kinases

All the 36,995 annotated proteins for the Pvu genome (v.2.1) were downloaded and scanned for the presence of putative kinase domains, as per the typical HMMs of the kinase domains (PF00069 and/or PF07714). In this first step, 1,800 proteins were found with significant alignments against these domains. From this set of alignments, 541 proteins were discarded for representing isoforms and 56 for not having a coverage of at least 50% of the corresponding kinase domain. These 56 sequences are likely related to atypical kinases or pseudogenes (Lehti-Shiu & Shiu, 2012; Liu et al., 2015). Out of the remaining 1,203 putative kinases, 775 returned from search criteria of PF00069, and 440 from search criteria of PF07714, with 6 PKs showing both domains (Supplementary Table S2).

The 1,203 PKs found were classified into 20 groups and 119 subfamilies through comparative alignments using HMMER and HMMs constructed with sequences from subfamilies of 25 other plant species (Lehti-Shiu & Shiu, 2012). In total, 1,197 PKs were confirmed by phylogeny (Supplementary Figs. S1-2; Supplementary Table S3). The group with the highest quantity of PKs was RLK-Pelle (∼70% of the amount of PKs), followed by CAMK (∼7%), CMGC (∼6%), TKL (∼5%), and STE (∼4%). Among the predicted kinases, six were considered to belong to an additional group called ‘Unknown’, which may represent specific subfamilies of Pvu. The distribution of PKs per subfamily had a mean of ∼10 (Supplementary Table S4) with a high dispersion (standard deviation of ∼17), caused by the presence of a few very large subfamilies. Among the RLK group, the RLK-Pelle_DLSV subfamily stood out as the most numerous (140), representing 11.6% of the total PKs of the species. In fact, the RLK-Pelle_DLSV subfamily was also the most numerous one in almost all 26 species analyzed, except in Smo (Supplementary Fig. S3). The closest species to Pvu regarding subfamilies’ composition were Ppe, Vvi and Mtr.

### 3.2. Kinase gene mapping and structural characterization

After identifying the PKs and classifying them into families and subfamilies, the genomic annotation information was used to position each PK gene along the Pvu genome. As a result, 1,191 PKs could be mapped to chromosomes, while 12 were located in scaffolds. The distribution of PKs per chromosome was extracted via GFF correspondences together with the measurement of intron counts in the related genes (Fig. 1A; Supplementary Table S5). There was not a noticeable concentration of the 1,991 PKs in any specific chromosome (Supplementary Table S6), and each of the remaining 12 PKs was located in a different scaffold. The greatest quantity of PKs was observed in chromosome 8 (172, 14.30%), and the least, in chromosome 10 (65, 5.40%). Although the largest chromosome contained the highest number of annotated PKs (chromosome 8 with ∼63 million base pairs (Mb) in length); the opposite was not observed in the shortest one (chromosome 6 with ∼31 Mb had 101 PKs estimated, while chromosome 10 had only 65 with a length of ∼44 Mb).

**Fig. 1.**
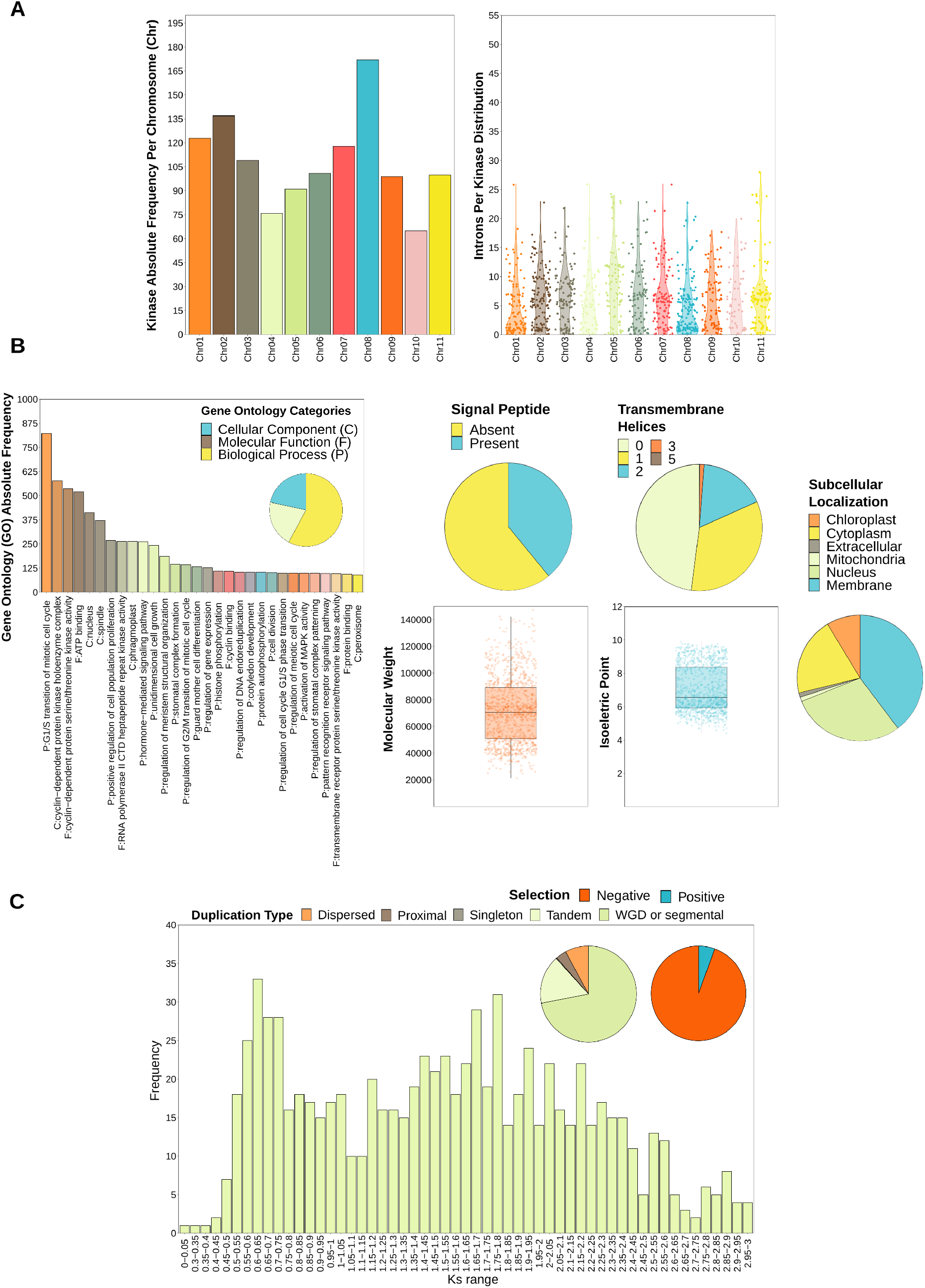
Descriptive analysis of kinase characteristics in *Phaseolus vulgaris* : (A) chromosomal distribution and intron occurrence; (B) presence of signal peptides and transmembrane helices, and distribution of molecular weights, isoelectric points (pIs), Gene Ontology (GO) terms, and subcellular localizations; and (C) duplication events.

We found that 163 PKs (13.5%) did not show introns in their gene structure. Most genes (835 or 69.4%) had up to 10 introns, while 182 PKs had between 11-20 introns (15.1%). For 23 genes (1.9%), more than 20 introns were predicted. In our study, we found 5.74 introns per kinase on average (median of 5), and the largest quantities observed were 28 (found in a member of PEK GCN2 subfamily), 26 (RLK-Pelle_LRR-XIIIb, RLK-Pelle_LRR-XIIIb, RLK-Pelle_LRR-XIIIb), and 24 (RLK-Pelle_DLSV) (Supplementary Table S5).

### 3.3. Protein kinase properties

In order to further characterize common bean PKs, we checked for the presence of additional protein domains with the HMMER and the Pfam database (Supplementary Table S7). Of the PKs analyzed, 563 showed only kinase-like domains, while for the remaining 640, 57 additional domains were noted (Supplementary Table S8). Some of these domains have relevant annotations indicating important functional potentialities. The five most prominent domains were Leucine rich repeat N-terminal domain 2 (LRRNT 2), Leucine rich repeat 8 (LRR 8), LRR 1, D-mannose binding lectin (B_lectin) and S-locus glycoprotein domain.

The vast majority of Pvu PKs (1,167, 97%) presented only one kinase domain, while 34 and two PKs contained two and three of such domains, respectively (Supplementary Table S9). These 36 PKs are distributed among 12 subfamilies. It is noteworthy that the subfamilies in which two or three kinase domains were found were, in order of abundance: AGC_RSK-2 (21), RLK-Pelle_RLCK-XI (3), RLK-Pelle_L-LEC (2), RLK-Pelle_WAK LRK10L-1 (2), RLK-Pelle_DLSV (1), RLK-Pelle_LRK10L-2 (1), RLK-Pelle_LRR-VIII-1 (1), RLK-Pelle_PERK-2 (1), CMGC_CDK-CCRK (1), CMGC_SRPK (1), AGC_NDR (1), and Group-PI-2 (1). Several other domains could also be found, providing increased degrees of complexity for the analyzed proteins (Supplementary Table S8). Up to 14 domains were predicted in the Phvul.005G025000.1.p protein, including the zf-RING_UBOX, Ank_2 and SH3_15 domains, in addition to Pkinase. The diversity of distinct domains observed (57), as well as their combinations, is extensive.

We could not obtain a consistent prognosis of PK subcellular localizations by all the selected software (WoLF PSORT, CELLO and LOCALIZER); therefore, we only considered predictions for PKs with a coincidence by at least two tools. Employing this approach, 697 PKs (∼60%) could have their localization predicted into six categories: chloroplast, cytoplasm, extracellular, mitochondria, nucleus, or membrane regions. The most prominent localizations were the membrane, cytoplasm and nucleus, to which 41.7, 24.4 and 17.5% of Pvu PKs were attributed, respectively (Supplementary Table S10; Fig. 1B).

The other protein properties evaluated were the pI, molecular weight, and presence of signal peptides and transmembrane helices (Supplementary Table S10; Fig. 1B). We found that ∼39% of Pvu PKs had an estimated presence of signal peptides. Transmembrane helices were found in ∼52.04% of PKs, separated in proteins with one (33.75%), two (17.04%), three (1.16%), and five helices (0.08%). Regarding pIs, the values found ranged from 4.42 to 9.9, with an average of 7.03 and a median of 6.56. Molecular weight values ranged from 21,379.91 to 181,740.93 kDa, illustrating the diversity of sizes of macromolecules, with 72,132.36 and 70,700.84 for mean and median, respectively.

We also performed a full GO annotation of Pvu PKs (Supplementary Table S11), which returned 19,061 different terms separated into biological process (∼58%), molecular function (∼21%) and cellular component (∼22%). The top 30 terms are presented in Fig. 1B and, for an easier interpretation of the results, a treemap containing all the GO terms related to biological processes was constructed with the REViGO tool (Supplementary Fig. S4). We could observe a clear prominence of terms related to the regulation of defense response, protein autophosphorylation, and post embryonic development.

Regarding the structural diversity and protein properties among PKs, we could observe distinct features between subfamilies (Supplementary Tables S12-S13). Although our analyses of Pvu PK genes did not reveal any clear distribution pattern in the intron quantity per kinase (Fig. 1A), it was possible to note that members of the same subfamily tended to have a similar number of predicted introns. For instance, all five members of the RLK-Pelle_LRR-VII-1 subfamily had only one intron, and all five members of RLK-Pelle_LRR-IV had three introns. Of the 118 subfamilies, 15 had members with the same number of introns, and for the remaining, most had a relatively conserved number of introns among their members. To get an overview of the number of introns of proteins in the same subfamily, the variance in each of them was analysed. We observed that 24 subfamilies presented only one member and, of the remaining 94 subfamilies, 15 had members with the same number of introns, i.e, with a variance equal to zero. Only six subfamilies showed variance above 10, indicating that members vary significantly in relation to the number of introns.

In addition to have the largest amount of PKs, we observed that RLK-Pelle_DLSV presented the most diverse set of domains (10 additional domains) and also the highest quantity of signal peptides, indicating a significant diversity of this family. Regarding the quantity of domains found in PKs, RLK-Pelle_LRR-III and RLK-Pelle_LRR-VII-1 followed RLK-Pelle_DLSV, presenting 5 and 4 additional domains respectively (Supplementary Table S13). The highest pI mean was observed in CK1_CK1 subfamily (9.61), followed by Group-Pl-4 (9.52) and RLK-Pelle_RLCK-IV (9.43). Interestingly, CMGC_Pl-Tthe subfamily presented the maximum molecular weight predicted with only one member.

### 3.4. Duplication analysis

From the investigation of PK origins through duplication events, we could find estimates for 1,167 PKs corresponding to 97% of the total kinome (Supplementary Table S14). The prominent origin was caused by WGD or segmental duplications with 839 PKs, followed by tandem (191), dispersed (92), and proximal (42) duplications. 3 PKs were singleton. Regarding collinearity events, Ka/Ks ratios ranged from 0.046 to 4.574, with an average of 0.397 (Supplementary Table S15; Supplementary Fig. S5). This ratio is used to estimate the balance between neutral mutations, purifying selection and beneficial mutations on a set of genes encoding homologous proteins. The calculation is based on the ratio between the number of non-synonymous substitutions per non-synonymous site in a given period of time and the number of synonymous substitutions per synonymous site, in the same period. In short, values above 1 for this equation are evidence of advantageous mutations; values below 1 imply pressure against change; and values close to 1 correspond to neutral effects over the period. However, positive and negative changes can cancel each other out over time. As we observe through PKs, there are cases of positive selection of substitutions, but the vast majority of changes, whose average was 0.397, seems to act against selection. Based on clock-like Ks rates, we also estimated the time at which these duplications occurred – which ranged from 1.2 to 229.1 million years ago (MYA) (Supplementary Table S15).

Tandem duplications were observed in 66 subfamilies (Supplementary Table S12), with the largest number of occurences in members of the RLK-Pelle group (22 in RLK-Pelle_DLSV, 19 in RLK-Pelle_LRK10L-2, 17 in RLK-Pelle_CrRLK1L-1, 8 in RLK-Pelle_LRR-III, and 7 in RLK-Pelle_WAK_LRK10L-1). By evaluating the distribution of GO terms in such tandemly duplicated PKs (Fig. 2A), we observed a similar profile to that observed in the total kinome (Supplementary Fig. S4), with the prominent terms related to response to stress.

**Fig. 2.**
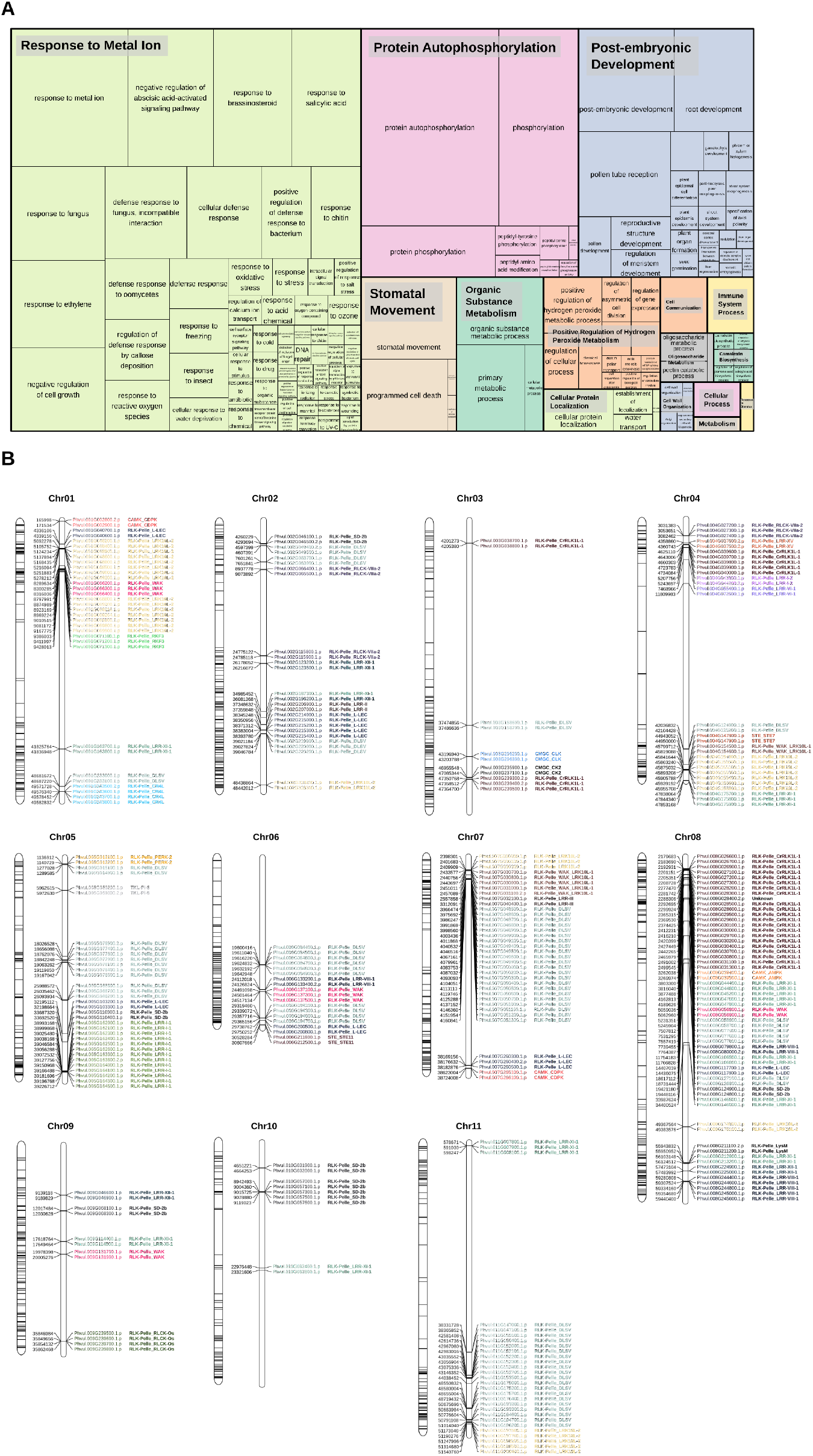
(A) Gene Ontology (GO) categories (biological processes) related to tandemly duplicated kinases. The size of the subdivisions within the blocks represents the abundance of that category in this set of kinases. The colors are related to the similarity to a representative GO annotation for the group. (B) Kinase distribution along chromosomes. For each chromosome, all genes with kinase domains are indicated on the left, and only the tandemly organized kinases are indicated on the right, colored and labeled according to the subfamily classification.

### 3.5. Gene expression and co-expression networks

In order to measure the expression level of each of the 1,203 PKs identified in this study in a broad range of conditions, data from transcriptome studies involving several genotypes and specific designs were obtained. Initially, we estimated the TPM values associated with each PK (Supplementary Table S16), and combined such quantifications per subfamily (Supplementary Table S17), averaging replicates and calculating a single value for each combination of control/non-control conditions, genotypes and tissues (Supplementary Table S18). From the heatmap constructed for the visualization of such quantifications (Fig. 3), we could observe grouping profiles according to the genotypes/tissues and experimental conditions, although with several overlays, indicating the complex expression underlying kinase subfamilies.

**Fig. 3.**
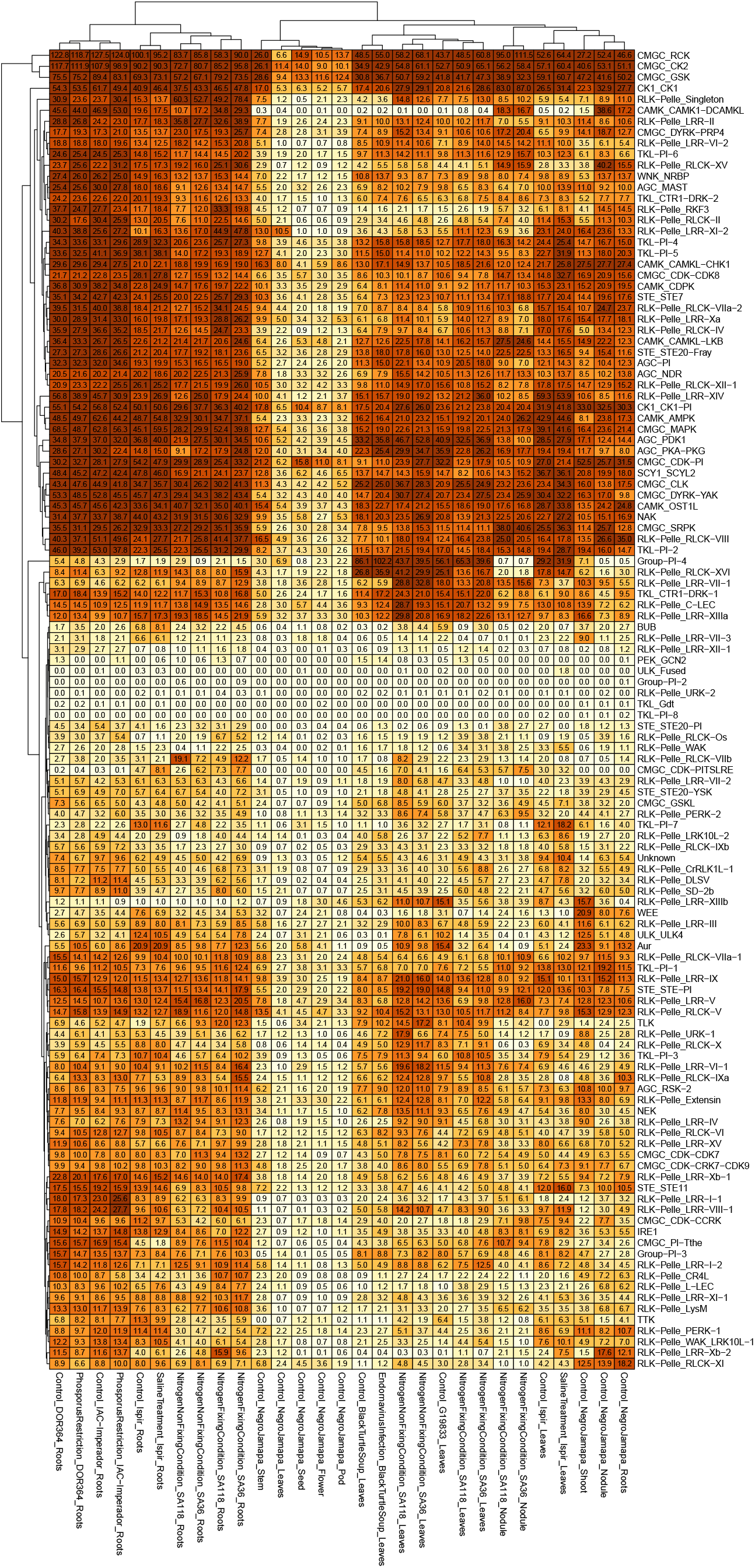
RNA expression profiles of *Phaseolus vulgaris* kinases, shown on a heatmap indicating the average sample values of different combinations of genotypes and tissues (columns) and considering the organization of kinase subfamilies (rows).

The top 5 mean expression values found were in CMGC_RCK, CMGC_CK2, CMGC_GSK, AGC_PDK1, and CK1_CK1-Pl subfamilies (Supplementary Table S19), which also presented the highest median measures. Regarding the maximum TPM values over samples, CMGC_RCK, CMGC_CK2, Group-Pl-4, CMGC_GSK, and CK1_CK1 represent the highest measures. CK1_CK1 presented the 6th highest expression, and, interestingly, although Group-Pl-4 did not present expressive expression values (99th highest expression), it was among the top 5 subfamilies with the largest variation of expression within samples. Other subfamilies with increased variation coefficients for the expression values within samples were Group-Pl-2, ULK_Fused, TKL-Pl-8, and CAMK_CAMK1-DCAMKL. It is noteworthy that TKL-Pl-8, ULK Fused, and Group-Pl-2 presented the lowest values for mean/median expression. By taking the lowest variation coefficients, the subfamilies with the most uniform expression across samples were RLK-Pelle_RLCK-V, AGC_RSK-2, RLK-Pelle_LRR-IX, CAMK_CAMKL-CHK1, and RLK-Pelle_Extensin.

In order to evaluate putative associations of the subfamilies’ expression with the profile of duplications and the quantity of PKs per subfamily, we performed correlations between such measures and the subfamilies’ TPMs for each combination of control/non-control conditions, genotypes and tissues (Supplementary Table S20). No significant Spearman correlation coefficients were found, being the largest values around 0.18, indicating that such an association is composed of joint factors which could not be easily captured by the measures evaluated.

Regarding the differences on the expression profile of control samples and samples under adverse conditions, we could infer an overall difference between such sets by using the heatmaps constructed (Supplementary Figs. S6-S7). However, as we employed samples from different studies, we performed a comparative analysis of such differences in terms of gene co-expression patterns rather than statistical tests (Fig. 4). In that sense, we modeled two different networks, one for control samples (Fig. 4A; Supplementary Fig. S8) and another one for samples under adverse conditions (Fig. 4C; Supplementary Fig. S9).

**Fig. 4.**
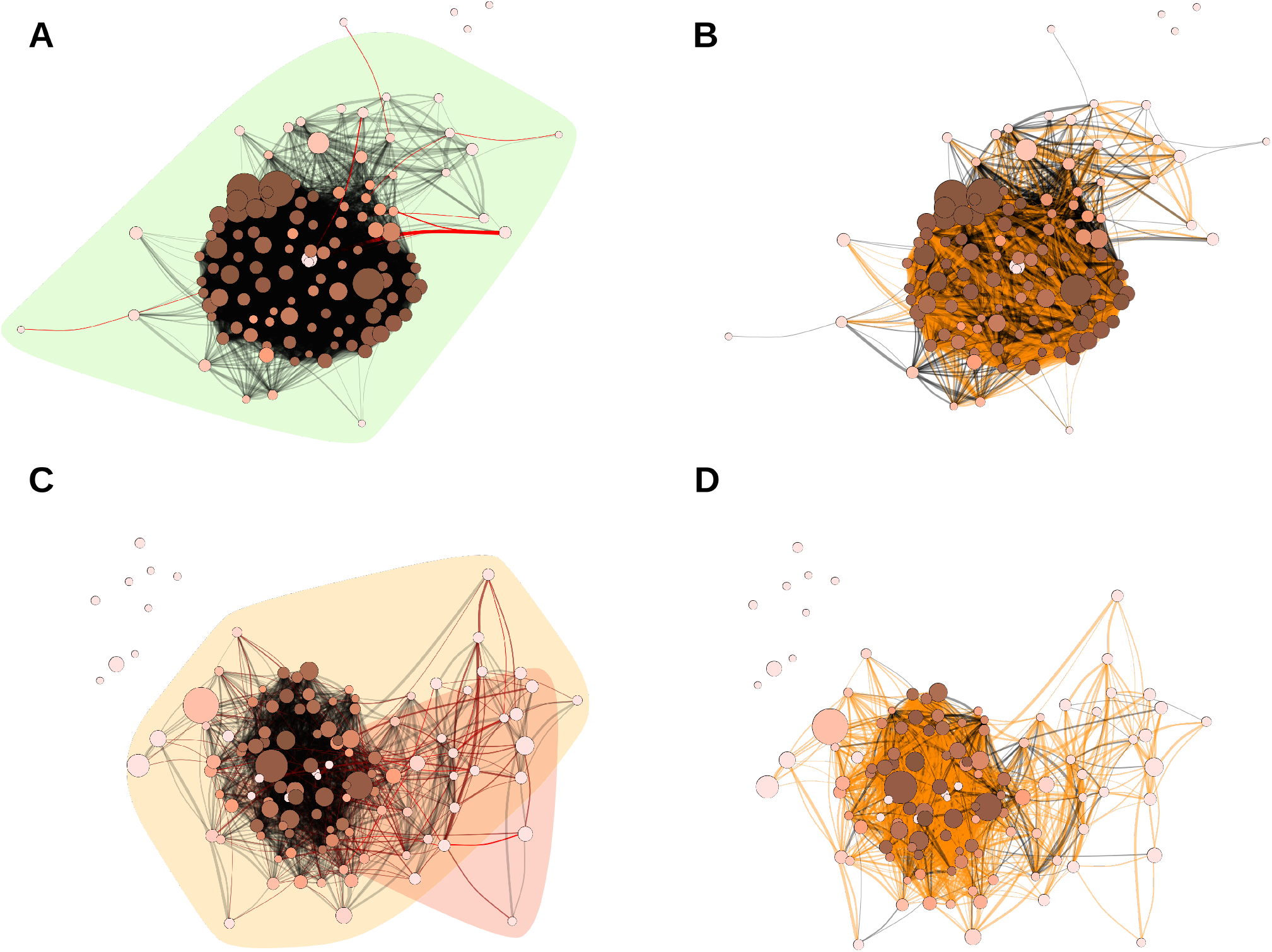
Coexpression networks for *Phaseolus vulgaris* (Phvul) kinase subfamilies. Each node corresponds to a different subfamily, its size corresponds to the average expression value for all kinases within the subfamily in different samples, and its color corresponds to the hub score and ranges from beige to dark brown. Each edge corresponds to a correlation with a Pearson correlation coefficient of at least 0.6. The correlation strength is represented by the edge’s width and the edge betweenness score is represented by the color (ranging from black to red, with red representing the highest values). (A) Phvul network (control samples) with the background colored according to the community detection analysis. (C) Phvul network (stress submitted samples) with the background colored according to the community detection analysis. (B) Phvul network (control samples) indicating the similarities with the Phvul network (stress submitted samples) in orange. (D) Phvul network (stress submitted samples) indicating the similarities with the Phvul network (control samples) in orange.

Although there was a common core structure between the networks modeled (Fig. 4B; Fig. 4B), several differences were identified. Firstly, we evaluated the presence of communities within the networks, and in contrast to a single member in the control network, the other one presented two different communities, one of them clearly separated from the main group. This indicates a more cohesive structure in the control network when compared to more sparse connections in the network affected by stress-related factors. In addition, hub and betweenness centrality measures were investigated for each one of the networks and clear distinctions could be pointed out.

As expected, in the control network, the hub scores for each kinase subfamily were bigger (Supplementary Table S21), standing out the PK subfamilies CMGC_CK2, CK1_CK1-Pl, TKL-Pl-4, CMGC_GSK, and STE_STE20-Fray. Concerning betweenness scores, the most vulnerable connections were those between the pairs of subfamilies CMGC_CDK-PITSLRE/RLK-Pelle_RLCK-X, Group-Pl-4/RLK-Pelle_RLCK-XVI, RLK-Pelle_LRR-VII-1/RLK-Pelle_LRR-XIIIb, and RLK-Pelle_RLCK-XI/TKL-Pl-8 (Supplementary Table S22). In the network with the samples under adverse conditions, on the other hand, more sparse hub scores were found (Supplementary Table S23), with the top 5 being CAMK_CDPK, CK1_CK1-Pl, TKL-Pl-2, TKL-Pl-4, and AGC MAST. Regarding betweenness measures, largest values were identified in this network contrasted to the control one, standing out the connections between the pairs RLK-Pelle_LRR-IX/RLK-Pelle_LRR-XV, RLK-Pelle_LRR-VII-1/RLK-Pelle_RLCK-X, AGC_RSK-2/ULK_ULK4, NEK/RLK-Pelle_RLCK-X, AGC_PKA-PKG/STE_STE-Pl (Supplementary Table S24).

## 4. Discussion

The number of PKs predicted for common bean (1,203) represents 3.25% of all predicted proteins for this species (36,995), an indicator of the importance of this superfamily. These results are similar to the percentage of PK genes in the genome of several other plants, such as 3.8% in maize (Wei et al., 2014), 3.4% in *A. thaliana* (Zulawski et al., 2014), 3.7% in grapevine (Zhu et al., 2018b). These number are, however, slightly inferior to the those found for the two closest Pvu relatives with kinomes compiled: cowpea and soybean, for which 4.3 and 4.7% of proteins were predicted as PKs, respectively (Ferreira-Neto et al., 2021; Liu et al., 2015). The methodology adopted by most of the studies mentioned above was the same – a HMM approach (Lehti-Shiu & Shiu, 2012) – allowing comparative inferences to be made between them. To enable inferences and comparisons with kinomes from other species, the criteria established for this work were similar to other studies on this subject (Aono et al., 2021; Liu et al., 2020, 2015; Singh et al., 2014; Wei et al., 2014; Zhu et al., 2018a,b; Zulawski et al., 2014). The high representativeness of the RLK-Pelle group among all kinases was noteworthy (Fig. 1). This occurrence is not surprising, as the high proportion of this group in the kinome of plants is unanimous; on average, RLK-Pelle PKs represent 68.5% of RLKs in all kinomes studied to date (Aono et al., 2021; Ferreira-Neto et al., 2021; Liu et al., 2020, 2015; Singh et al., 2014; Wei & Li, 2019; Wei et al., 2014; Yan et al., 2018; Zhu et al., 2018a,b; Zulawski et al., 2014). Members of the RLK/Pelle family are directly involved in plant development, defense against pathogens, and responses to abiotic stresses (Lehti-Shiu & Shiu, 2012). The evolution of plants is likely associated with the expansion of subfamilies of this group, with special regard to the perception of pathogen signals and the subsequent triggering of immune responses. In fact, studies have shown an association between molecular markers, genes encoding RLK-LRR proteins and disease resistance (Binagwa et al., 2021; Vaz Bisneta & Gonçalves-Vidigal, 2020). The second most representative group among the kinases was the CAMK. Kinases of this group have been shown to act as primary sensors and to participate in various biological processes, such as the perception of calcium signals, the regulation of plant growth and development, and responses to biotic and abiotic stresses. According to Wei et al. (2014), the expansion of the CDPK family could be a consequence of the adaptive evolution of plants to perceive calcium signals. In our study 39 CAMK-CDPK proteins were found within this group.

Regarding the distribution of introns in common bean PKs, the maximum intron number observed was 28 – the same number found for soybean (Liu et al., 2015) and cowpea (Ferreira-Neto et al., 2021). Among available kinomes, the highest numbers of introns were found in grapevine (49) (Zhu et al., 2018b), sugarcane (52) (Aono et al., 2021), and pineapple (67) (Zhu et al., 2018a). The mean introns number found for common bean PKs (5.74) is lower than those found for strawberry (Liu et al., 2020) and pineapple (Zhu et al., 2018a), which were 6.45 and 6.59, respectively. Of the 118 subfamilies of common bean PKs, 15 had members with the same number of introns, and for the remaining, most had a relatively conserved number of introns among their members, as was also observed for soybean (Liu et al., 2015). Additionally, 163 common bean PK genes (13.5%) did present introns. In wheat, 11.9% of PKs showed no introns in their gene structure (Wei & Li, 2019), 9.5% in pineapple (Zhu et al., 2018a). Soybean, cowpea and grapevine have 12.1, 13.6 and 16.6%, respectively (Ferreira-Neto et al., 2021; Liu et al., 2015; Zhu et al., 2018b). In wheat, only 13.91% of PKs have more than 10 introns (Wei & Li, 2019).

At the family level, there is evidence of a link between the structural diversity of genes that are members of gene families and their evolution (Wei et al., 2014). In our study, the variation in the number of introns among PKs within the same subfamily was not large. In general, subfamilies showed conserved exon-intron structures, as observed by Yan et al. (2017), which may be related to their phylogenetic relationship. Most maize PK genes clustered in the same subfamily share similar intron structure, suggesting that intron gain and loss events contribute to the structural evolution of families (Wei et al., 2014). Liu et al. (2015) compared their results in soybean with those obtained for rice and maize, noting great similarity, referring the evolutionary history of PKs to times prior to the evolution of mono- and dicotyledons. Our results for common bean are similar to those for soybean, corroborating these conjectures. Divergent gene structures in different phylogenetic subfamilies may represent the expansion of the gene family (Wei et al., 2014), with kinase families having their own evolutionary expansions from the point of divergence (Liu et al., 2015). Conservation in the exon-intron structure of PKs, associated with growth and development processes, may originate from the emergence of land plants and thus be perpetuated (Yan et al., 2017).

### 4.1. Kinase protein properties

The distribution of kinase domains found for common bean was quite similar to that observed for sorghum (Aono et al., 2021), grapevine (Zhu et al., 2018b), wheat Yan et al. (2017), and soybean (Liu et al., 2015). Regarding the number of proteins with multiple kinase domains, there was a variation in the number of subfamilies and members; the subfamilies that contained the most multi-kinases members were AGC_RSK-2 and RLK-Pelle_RLCK-XI, as equally noted for soybean (Liu et al., 2015). Additionally, in sorghum, sugarcane (Aono et al., 2021), grapevine (Zhu et al., 2018b), pineapple (Zhu et al., 2018a), and wheat (Yan et al., 2017), the AGC_RSK-2 subfamily was also the most numerous. The second most numerous families were found to be RLK-Pelle_WAK in sorghum (Aono et al., 2021), AGC NDR in wheat (Yan et al., 2017), and RLK-Pelle_DLSV in sugarcane (Aono et al., 2021), grapevine (Zhu et al., 2018b) and pineapple (Zhu et al., 2018a). Only 36 (3%) of common bean PKs presented more than one kinase domain, which were distributed into 16 families. In soybean, the 74 PKs with such characteristics were distributed between 18 subfamilies, the most numerous being AGC_RSK-2 (38) and RLK-Pelle_RLCK-XI (7) (Liu et al., 2015). In sugarcane, the 228 proteins with multiple kinase domains are distributed into 49 subfamilies, the most numerous being AGC_RSK-2 (50) and RLK-Pelle_DLSV (29), while in sorghum the 49 proteins are distributed into 13 subfamilies, with AGC_RSK-2 (19) and RLK-Pelle_WAK (11) being the most numerous (Aono et al., 2021). Differently, in strawberry, of the 954 PKs analyzed, 920 presented two or more kinase domains and, therefore, 34 presented only one kinase domain (Liu et al., 2020). In cowpea only 6 PKs have only 1 kinase domain, while the rest have a higher number of kinases (Ferreira-Neto et al., 2021).

The importance of predicting the subcellular localization of each one of the proteins of a species lies in determining its place of action, which can in turn suggest its function (Zhu et al., 2018a) in association with further information, such as structural domains. The fact that many common bean PKs are located in the cell membrane suggests the relevance of this superfamily in perceiving the extracellular environment and transducing vital information into cells (Zhu et al., 2018a). In sorghum, sugarcane (Aono et al., 2021), cowpea (Ferreira-Neto et al., 2021), wheat (Wei & Li, 2019), pineapple (Zhu et al., 2018a), grapevine (Zhu et al., 2018b), soybean (Liu et al., 2015), and *A. thaliana* (Zulawski et al., 2014) the PKs predicted to locate at the cell membrane are also the majority, with percentages ranging from 27.42% in grapevine to 49.63% in soybean. In strawberry, on the other hand, 55.77% of PKs were predicted to locate at the nucleus (Liu et al., 2020). In our study, 501 PKs had their subcellular localization predicted to the plasma membrane and, among these, 486 (97%) were classified as RLK-Pelle – reinforcing the importance of these proteins in cell signaling. While the vast majority of membrane PKs are RLKs, it cannot be said that all RLKs are membrane PKs. While most (58%) of these proteins were predicted to locate at the membrane, 13.7% of RLK-Pelle proteins predicted to be cytoplasmic, 9.8% extracellular and 9.5% nuclear; additionally, 4.5 and 4.4% of these proteins were predicted to locate at chloroplasts and mitochondria, respectively. The observations made for common bean were very similar to those obtained for soybean, including the location of the RLKs, which were also essentially located at the membrane (Liu et al., 2015). In strawberry, on the other hand, only 45.4% of the RLKs were predicted to locate at the membrane (Liu et al., 2020). In pineapple, 38% of PKs were predicted to be located at the membrane and more than half of RLKs were membrane-located (Zhu et al., 2018a). PKs have great importance in sensing the environment and its response at the gene-expression level. The results observed for cowpea (Ferreira-Neto et al., 2021) are similar to the results found in our study.

Regarding PK pIs, the results found for common bean were similar to those of other species, such as sorghum, sugarcane (Aono et al., 2021), grapevine (Zhu et al., 2018b), and especially cowpea (Ferreira-Neto et al., 2021). However, for molecular mass, considerable differences were observed. The minimum molecular weight value of common bean PKs was higher than that observed for sugarcane, cowpea and grapevine, while the maximum value was lower than those found for sorghum, sugarcane, cowpea and grapevine. In grapevine, members of the same family share number of introns, pIs and molecular weight (Zhu et al., 2018b), while for cowpea the values of pI and molecular weight within families are highly variable (Ferreira-Neto et al., 2021). Our results follow the trend observed for cowpea, with highly variable values within the same family.

### 4.2. Duplication events

Alike other species (Aono et al., 2021; Ferreira-Neto et al., 2021; Liu et al., 2015; Zhu et al., 2018a,b), the Pvu kinome presented a high percentage of PK gene pairs with a Ka/Ks ratio below 1, indicating that they are under purifying selection. This indicates that selection has acted to conserve the structure and stabilize the function of PKs along their evolutionary history. In eukaryotes, this is thought to ensue an early phase of relaxed constraint or even near-neutrality for diversification (Lynch & Conery, 2000), and possibly occurred during PK evolution due to their vital importance in diverse biological processes (Janitza et al., 2012).

Both the presence of duplicated genes under purifying selection and the average Ka/Ks rate of common bean PKs (0.397) are concordant with previous findings from other gene families of this species, such as Dof (Ito et al., 2017), SBP transcription factors (Ilhan et al., 2018), CAMTA (Büyük et al., 2019), SRS (Büyük et al., 2022), and BURP domain-containing genes (Kavas et al., 2021). However, none of these studies – which analysed much smaller gene families – reported the high Ks values and distant dates of duplication we found for common bean PKs, dating up to 229 MYA. We observed a distinct peak in Ks values ranging around 0.65, corresponding to duplication events occurring 50 MYA; this coincides with a major WGD experienced by the Fabaceae, estimated to have taken place 58 MYA (Lavin et al., 2005). A second, less evident peak can be observed in Ks values around 1.5-1.7, corresponding to duplications from 116-130 MYA; this can be associated with a whole-genome triplication event that took place in the core eudicots lineage, pinpointed at 117 MYA (Jiao et al., 2012). It is very likely that these two polyploidization events represent major forces in the diversification of common bean PKs, as also reported for legume transcription factor repertoires (Moharana & Venancio, 2020). Oddly, the influence of neither of these events was detected in the kinome of cowpea, a close relative of common bean. In the kinome of the slightly more distantly-related soybean, the Fabaceae-specific WGD Ks peak can be observed – although it is overshadowed by duplications arising from this species’ more recent, lineage-specific, WGD ∼13-59 MYA (Liu et al., 2015; Schmutz et al., 2010).

Specific PK subfamilies had a more pronounced occurrence of tandem duplications, mostly from the RLK-Pelle group (RLK-Pelle_DLSV, RLK-Pelle_LRK10L-2, RLK-Pelle_CrRLK1L-1, RLK-Pelle_LRR-III, and RLK-Pelle_WAK LRK10L-1). In addition, RLK-Pelle_DLSV and RLK-Pelle_LRR-III were among the subfamilies with the largest diversity of protein domains. As tandemly duplicated PKs are known to be associated with stress responses (Freeling, 2009), we could also evidence the expansion of the scope of functionality of these subfamilies.

### 4.3. Gene expression estimation

In our study, we incorporated several RNA-Seq datasets for estimating Pvu kinome expression, which enabled a broad overview of the PK expression across different common bean genotypes and tissues. Although the most pronounced subfamilies in PK quantities were RLK-Pelle_DLSV (11.64%), RLK-Pelle_LRR-XI-1 (5.15%), RLK-Pelle_CrRLK1L-1 (4.32%), and RLK-Pelle_LRK10L-2 (4.41%), we found different subfamilies with the largest expression values (CMGC_RCK, CMGC_CK2, CMGC_GSK, AGC_PDK1, and CK1_CK1-Pl). Such finding indicates that even if a PK subfamily is highly abundant across the genome, its expression might not reflect it, as already pointed out by other kinome studies (Aono et al., 2021; Liu et al., 2015). Indeed, by evaluating the correlation between the PK abundance per subfamily and their expression, we did not find significant associations (Supplementary Table S20).

Different members of CMGC group presented the largest expression values, as also reported by Aono et al. (2021); Liu et al. (2015); Zhu et al. (2018b). This result reinforces the high conservation of this group across several plant species (Kannan & Neuwald, 2004), and its multiple functions with effects on several signalling mechanisms (Wrzaczek et al., 2007). Several subfamilies presented variable expression values across their representatives, as highlighted by the high variation coefficients calculated (Supplementary Table S19), which corroborates the specific activation of PKs (Zhu et al., 2018a). Interestingly, although Group-Pl-4 subfamily did not present expressive expression values (99th highest expression), it was among the top 5 subfamilies with the largest variation of expression within samples and also with one of the maximum expression values observed among the entire kinome in a stress associated sample. In addition to being highly conserved (Lehti-Shiu & Shiu, 2012), Zhu et al. (2018a) already reported the potential involvement of such a subfamily with photosynthesis.

Finally, modelling different coexpression networks made it possible the definition of several inferences across PK subfamilies interaction patterns, distinguished in two different structures for modelling control and stress related samples. The use of complex networks for modelling biological systems has enabled important contributions in the decipherment of unknown molecular associations across the literature (Fait et al., 2020; Tai et al., 2020; Zhang & Yin, 2020). In our study, each PK subfamily represents an element in the network (a node) and their putative associations (edges) are estimated through linear correlations, which indicate PK subfamilies that are functionally cohesive, co-regulated or correspond to similar pathways (Mitra et al., 2013). From such a structure, network measures can be used for biological inferences, including central elements in the network structure (hubs), which are generally associated with regulators over the biological mechanisms modeled (Barabasi & Oltvai, 2004), and also connections with elevated network vulnerability (edges with high betweenness), i.e. connections permeating a high flow of communication between network elements. Considering the PK networks modeled, edges with high betweenness measures may represent crucial mechanisms for the maintenance of the overall PK interactions (Aono et al., 2021).

Although possessing a common core structure, the networks modeled presented several differences in their topology. First, the detachment of the network with samples under adverse conditions into two communities potentially indicates the disturbance of the previous network because of external factors into the complex system, namely the different adverse circumstances in which the genotypes were evaluated. As already known, the activation of PKs is directly affected by external stimuli and stress factors (Jaggi, 2018; Morris, 2001), and this aspect can be inferred from the networks modeled. Additionally, the large quantity of connections in the control network indicates a more cohesive structure with less vulnerability points; this suggests that the subfamily interactions presented a more synergistic activity than the interactions between the expression of subfamilies under stress. Indeed, such finding can be also visualized in the connections with higher betweenness values (Fig. 4).

In the network of control samples, we only found points of vulnerability between single subfamilies and the main core group, formed by a cohesive set of PK subfamilies interactions. However, in the other network, such edges with high betweenness seem to have a bigger impact into the network architecture (Fig. 4C). Members of the subfamilies RLK-Pelle_RLCK and RLK-Pelle_LRR were present in the edges among the top 5 betweenness values of both networks. Interestingly, other subfamilies in the vulnerable edges of the control network (TKL-Pl-8, CMGC_CDK-PITSLRE, and Group-Pl-4) were disconnected elements in the stress network. This demonstrates that the existent vulnerabilities become more pronounced in adverse conditions, and also reinforces the importance of the RLK-Pelle group (Bolhassani et al., 2021). By contrasting the other subfamilies present in the high betweenness edges in the stress related network with their connection profile in the control network, we can visualize that AGC_RSK-2, AGC_PKA-PKG, and NEK presented median hub scores, i.e. they have a significant amount of connections, which are significantly reduced in the network modeled with samples under adverse conditions. In the same way, STE_STE-Pl subfamily presented the same profile, but with a more elevated hub score, which was close to the top values in the control network. Such findings corroborate the potential of biological inferences with the use of complex networks and highlight this set of PK subfamilies for deeper investigations over Pvu stress responses.

Regarding the key elements in both networks, measured through hub scores, we found CK1_CK1-Pl and TKL-Pl-4 among the top 5 in both structures. Although we found differences in other hub elements, similar connection profiles could be observed. For instance, CMGC_CK2 and CMGC_GSK ranked in the top 5 hubs in the control network, and in the network modeled with samples under adverse conditions such families did not present low hub scores. The same was observed for the other hubs of the adverse network (CAMK_CDPK, TKL-Pl-2, and AGC MAST). Even not being in the top 5 of the control network, the values were close to the highest hub score. Interestingly, the subfamily STE_STE20-Fray was considered a hub in the control network, however in the adverse related network it had a low hub score in the adverse condition, which shows a probable impact of stress into this PK subfamily.

## 5. Conclusion

The common bean has a large importance for agriculture, representing a good source of nutrition. Considering the well-established role of PKs over stress responses and the diverse stresses affecting bean production, the characterization performed in our study represents an important contribution to Pvu research, cataloging a vast and rich reservoir of data. By profiling 1,203 Pvu PKs, we provided significant insights into Pvu PK organization, highly variable functional profile, structural diversity and evolution, and expression patterns. Finally, by modelling the PK interactions through coexpression networks, we could highlight a set of PK subfamilies potentially associated with bean stress responses.

## Supporting information

Supplementary Results

## Acknowledgements

This work was supported by grants from the Fundação de Amparo à Pesquisa do Estado de São Paulo (FAPESP), the Fundação de Amparo à Pesquisa do Estado de Minas Gerais (FAPEMIG), the Conselho Nacional de Desenvolvimento Científico e Tecnológico (CNPQ), and the Coordenação de Aperfeiçoamento de Pessoal de Nível Superior (CAPES, Computational Biology Programme). AA received a PhD fellowship from FAPESP (2019/03232-6) and RP received a PhD fellowship from FAPESP (2019/21682-9).

## Author contributions

AA, RP and WP performed all analyses and wrote the manuscript. CD and FC assisted in the kinase functional analyses. WP, RK and AS conceived the project. All authors reviewed, read and approved the manuscript.

## Supplementary Tables

**Table S1**. Organization of bean RNA-Seq experiments.

**Table S2**. Kinase domain annotation of the 1,203 bean protein kinases.

**Table S3**. Subfamily kinase classification of the bean 1,203 kinases.

**Table S4**. Bean kinase subfamily quantification.

**Table S5**. Localization and intron quantity of the 1,203 bean kinases.

**Table S6**. Bean kinase distribution across chromosomes.

**Table S7**. Domain annotation of the 1,203 bean protein kinases.

**Table S8**. Domain organization of the 1,203 bean protein kinases.

**Table S9**. Kinase domain organization for proteins with multiple kinase domains.

**Table S10**. Compositional analyses of the 1,203 kinases.

**Table S11**. Gene Ontology (GO) annotations for the 1,203 bean kinases.

**Table S12**. Characteristics of bean kinase subfamilies.

**Table S13**. Presence of domains presence across bean kinase subfamilies.

**Table S14**. Duplication origin of the bean 1,203 kinases.

**Table S15**. Collinearity events and Ka/Ks values of bean protein kinases.

**Table S16**. Kinase TPM values across samples.

**Table S17**. Kinase subfamily quantification across samples.

**Table S18**. Kinase subfamily quantification across tissues in the selected genotypes.

**Table S19**. Descriptive statistics of subfamily expression across kinase subfamilies.

**Table S20**. Spearman correlation of average TPM values in bean genotypes/tissues with kinase subfamily quantities.

**Table S21**. Kinase subfamily coexpression network characterization (control samples).

**Table S22**. Edge betweenness values calculated across the bean coexpression network (control samples).

**Table S23**. Kinase subfamily coexpression network characterization (adverse samples).

**Table S24**. Edge betweenness values calculated across the bean coexpression network (adverse samples).

## Supplementary Figures

**Fig. S1**. Phylogenetic analysis of the identified protein kinases in *Phaseolus vulgaris* (Phvul). Each protein is separated on the right side of the tree and is presented with its classification with respect to the kinase subfamilies, which are colored to represent the differences among subfamilies.

**Fig. S2**. Phylogenetic analysis of the identified protein kinases in *Phaseolus vulgaris* in a circular layout. Each protein is colored with respect to the kinase subfamily classification.

**Fig. S3**. Kinase subfamily quantification analysis in different plant species. Each row indicates a different subfamily and each column a plant species, and the numbers of kinases are noted. This heatmap is colored according to the distribution of quantities present in the datasets on a scale of beige to dark red.

**Fig. S4**. Gene Ontology (GO) category annotation of biological processes in the entire set of *Phaseolus vulgaris* kinases. The size of the subdivisions within the blocks represents the abundance of that category in this set of kinases.

**Fig. S5**. Segmental duplication events in the *Phaseolus vulgaris* genome. The colors indicate the selection type of the gene pair duplication (gray indicates negative selection and orange positive selection).

**Fig. S6**. RNA expression profiles of *Phaseolus vulgaris* kinases (control samples), shown on a heatmap indicating the average sample values of different combinations of genotypes and tissues (columns) and considering the organization of kinase subfamilies (rows).

**Fig. S7**. RNA expression profiles of *Phaseolus vulgaris* kinases (stress submitted samples), shown on a heatmap indicating the average sample values of different combinations of genotypes and tissues (columns) and considering the organization of kinase subfamilies (rows).

**Fig. S8**. Coexpression networks for *Phaseolus vulgaris* kinase subfamilies (control samples). Each node corresponds to a different subfamily, its size corresponds to the average expression value for all kinases within the subfamily in different samples, and its color corresponds to the hub score and ranges from beige to dark brown. Each edge corresponds to a correlation with a Pearson correlation coefficient of at least 0.6. The correlation strength is represented by the edge’s width and the edge betweenness score is represented by the color (ranging from black to red, with red representing the highest values).

**Fig. S9**. Coexpression networks for *Phaseolus vulgaris* kinase subfamilies (stress submitted samples). Each node corresponds to a different subfamily, its size corresponds to the average expression value for all kinases within the subfamily in different samples, and its color corresponds to the hub score and ranges from beige to dark brown. Each edge corresponds to a correlation with a Pearson correlation coefficient of at least 0.6. The correlation strength is represented by the edge’s width and the edge betweenness score is represented by the color (ranging from black to red, with red representing the highest values).

