## Supplementary figures and images for "Genome-wide characterization of the common bean kinome: catalog and insights into expression patterns and genetic organization"

### FigS2.png

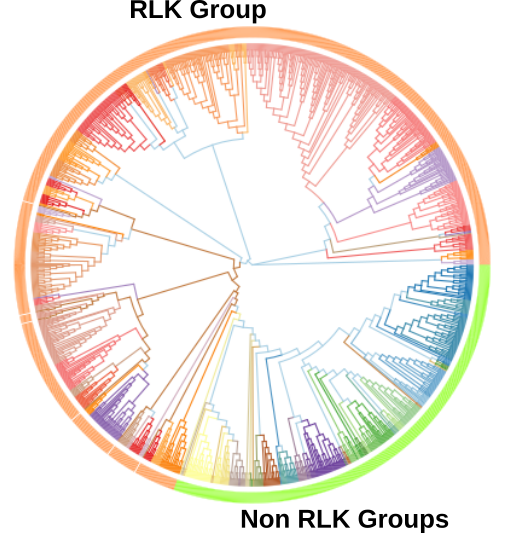

### FigS3.png

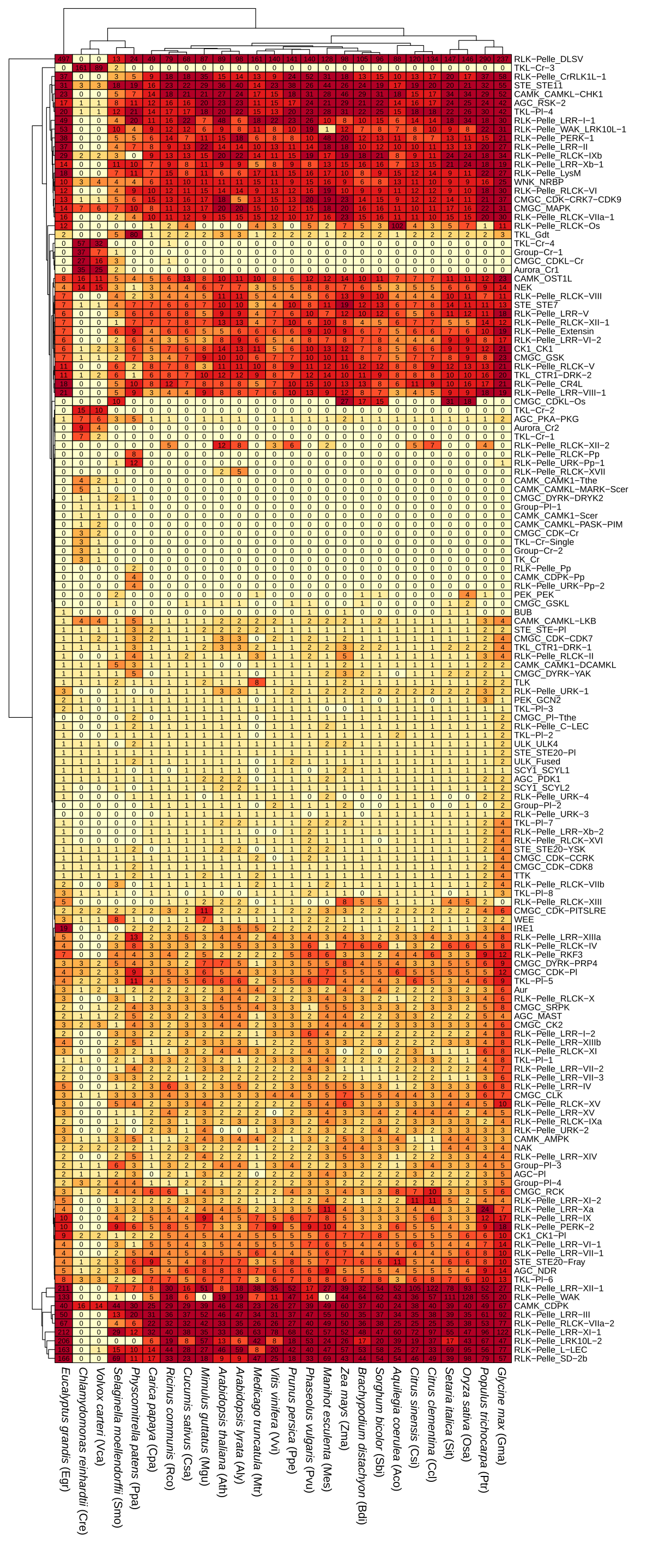

### FigS4.png

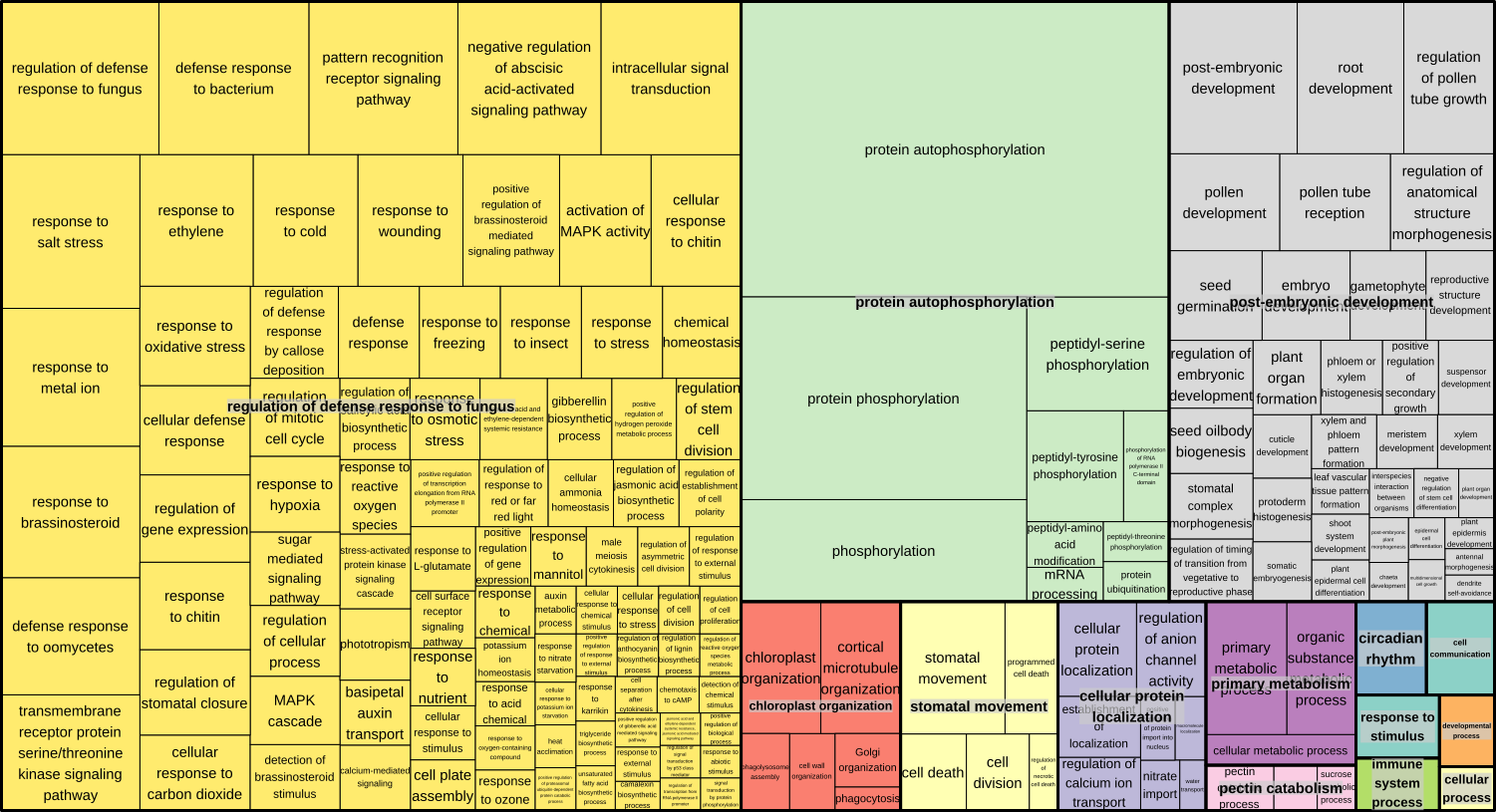

### FigS5.png

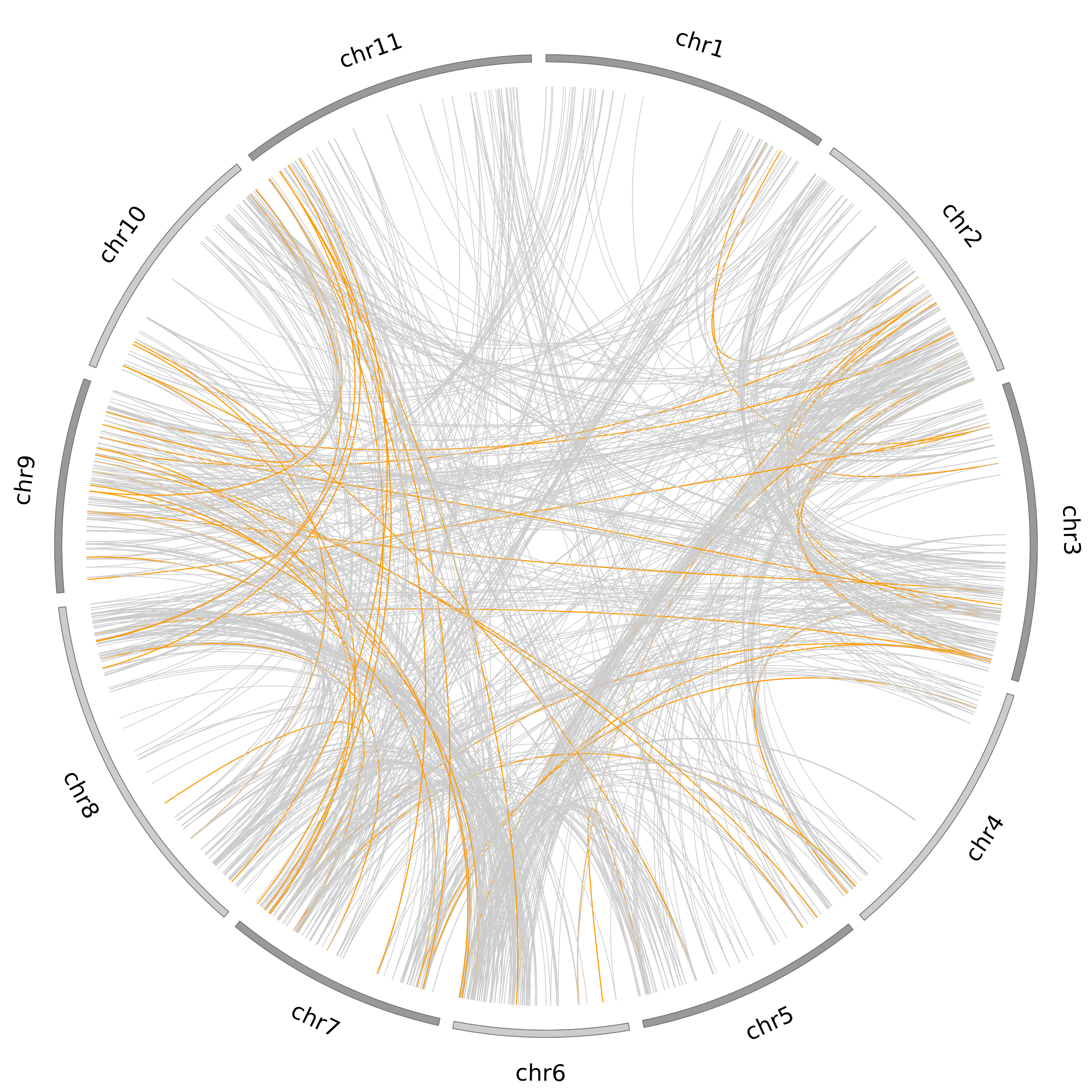

### FIgS6.pdf

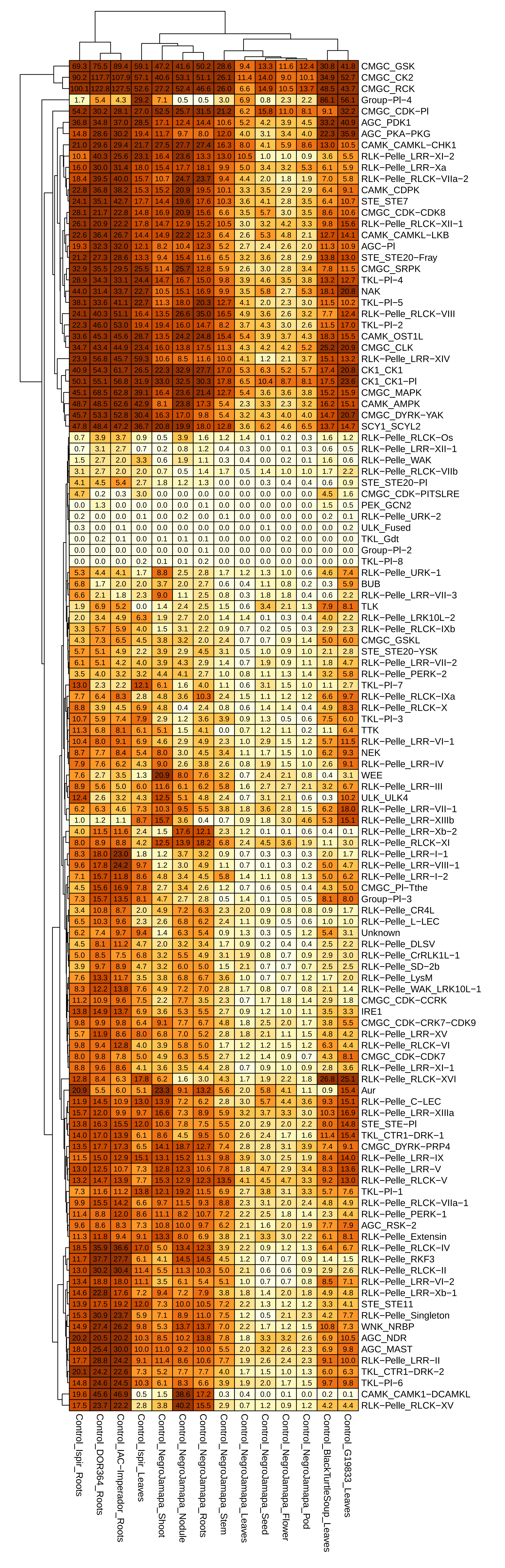

### FigS7.pdf

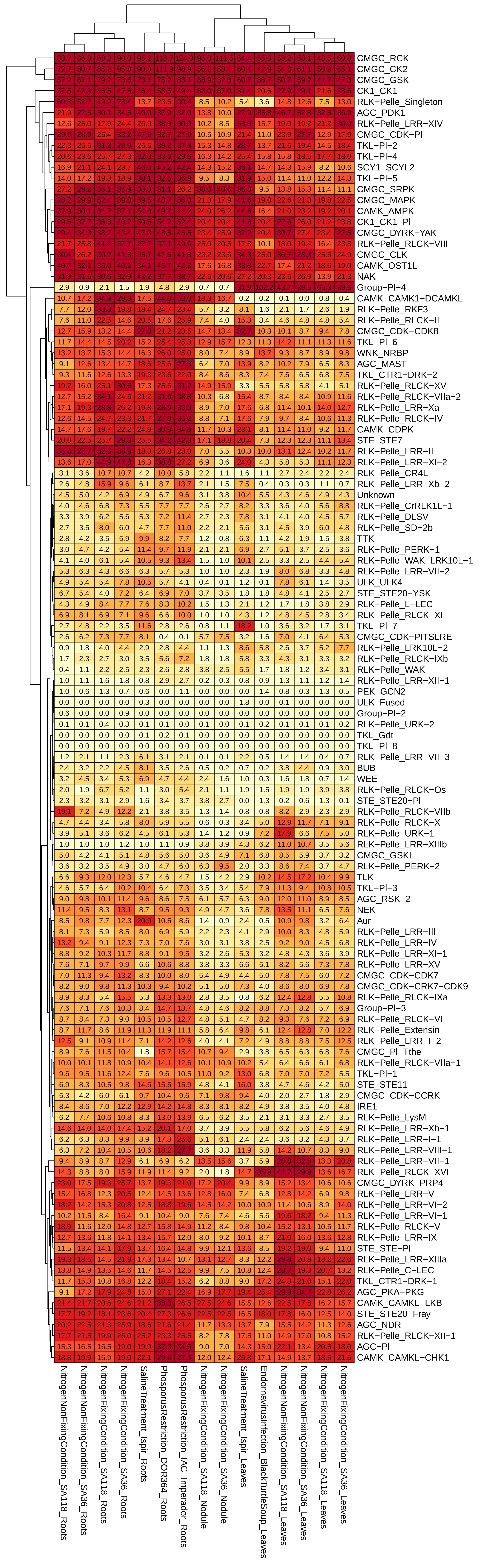

### FigS8.png

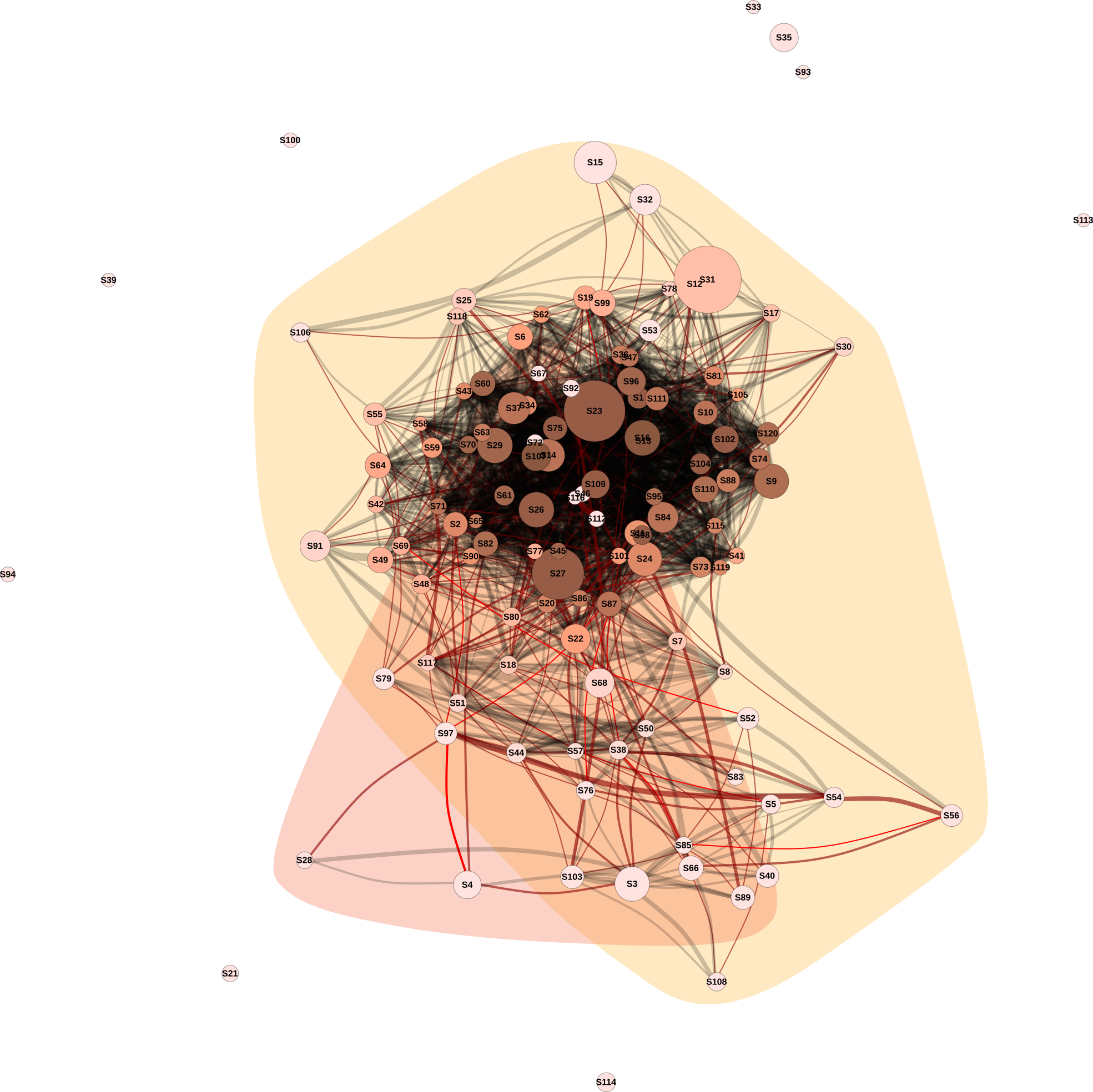

### FigS9.png

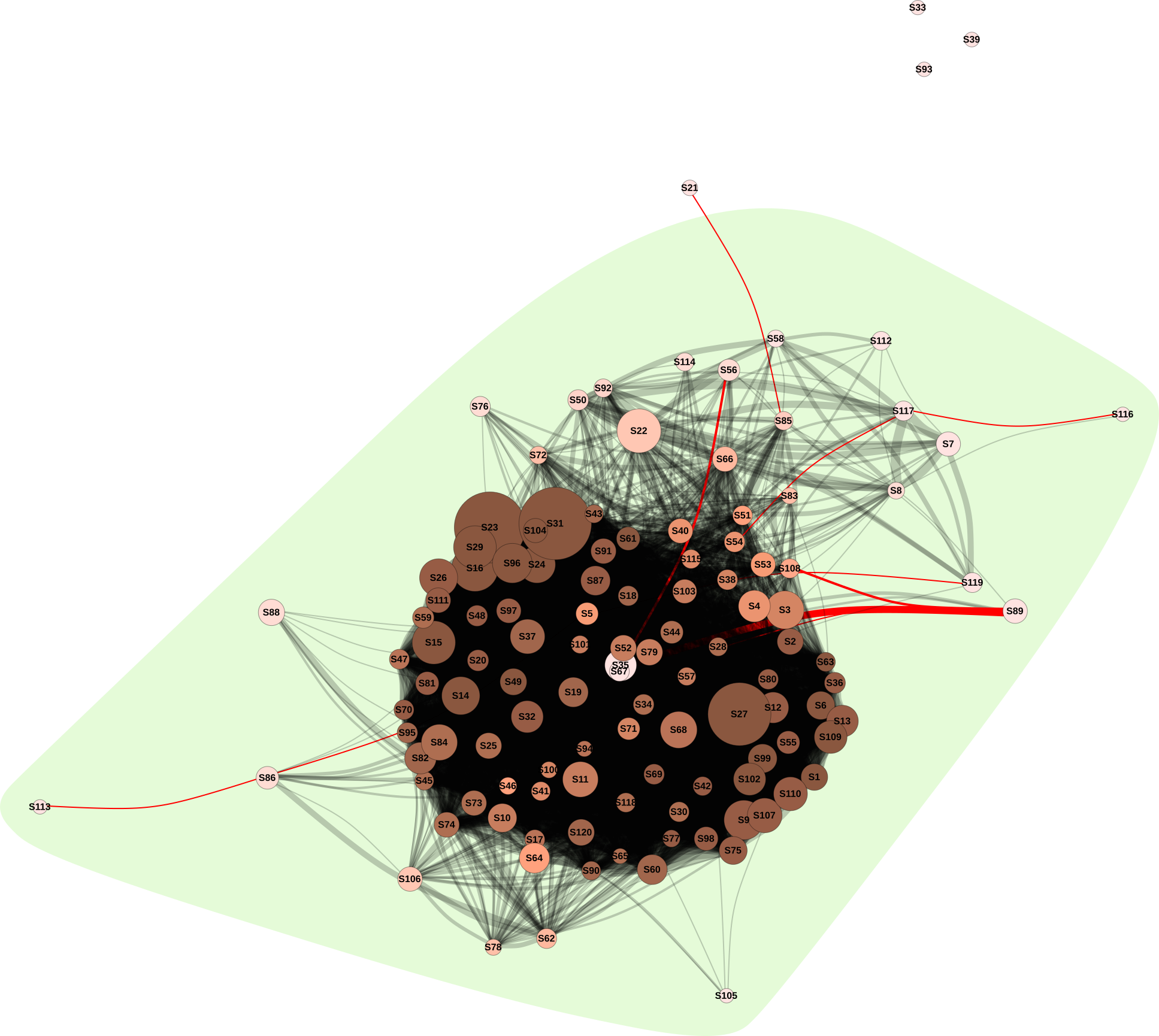
